# Dynamic oxygen-enhanced MRI of the lung at 3 T: feasibility, repeatability and reproducibility

**DOI:** 10.1101/2023.04.09.536144

**Authors:** Mina Kim, Josephine H. Naish, Sarah H. Needleman, Marta Tibiletti, Yohn Taylor, James P. B. O’Connor, Geoff J. M. Parker

**Affiliations:** Centre for Medical Image Computing (CMIC), Department of Medical Physics and Biomedical Engineering, University College London, London, United Kingdom; Bioxydyn Limited, Manchester, United Kingdom; BHF Manchester Centre for Heart and Lung Magnetic Resonance Research (MCMR), Manchester University NHS Foundation Trust, Manchester, United Kingdom; Division of Cancer Sciences, University of Manchester, Manchester, United Kingdom; Division of Radiotherapy and Imaging, Institute of Cancer Research, London, United Kingdom

**Keywords:** Lung, OE-MRI, 3 T, oxygen-enhanced MRI, dynamic, free-breathing, repeatability, reproducibility

## Abstract

**Purpose:** Dynamic T_1_-weighted lung oxygen enhanced MRI (OE-MRI) is challenging at 3 T due to decreased longitudinal relaxivity of oxygen and increased magnetic susceptibility difference between air and tissue interfaces relative to 1.5 T, leading to poor signal quality. In this work, we evaluate the robustness of an alternative T_2_*-sensitised lung dynamic OE-MRI protocol in humans at 3 T.

**Methods:** Simulations were performed to predict OE contrast behaviour and optimise the MRI protocol. Sixteen healthy subjects underwent dynamic free-breathing OE-MRI acquisitions using a dual echo RF-spoiled gradient echo acquisition at 3 T on two MRI scanners at different institutions. Non-linear registration and tissue density variation correction were applied. Percent signal enhancement (PSE) maps and ΔR_2_* were derived. Intra-class correlation coefficient (ICC) and Bland-Altman analyses were used to evaluate reproducibility of the OE indices across two sites and vendors as well as scan-rescan repeatability.

**Results:** Simulations and experimental data show negative contrast on oxygen inhalation due to substantial dominance of ΔR_2_* at TE longer than 0.2 ms when using our chosen flip angle and TR. Mean PSE values were TE dependent and mean ΔR_2_* was 0.14 ms^-1^ ± 0.03 ms^-1^, ICC values for intra-scanner (ICC_intra_) and inter-scanner (ICC_inter_) variability for OE indices were high (ICC_intra_ > 0.74; ICC_inter_ = 0.70) and 95% limits of agreement showed strong agreement of repeated measures.

**Conclusion:** Our results demonstrate excellent scan-rescan repeatability for the PSE indices and good reproducibility for ΔR_2_* across two sites and vendors, suggesting potential utility in multi-centre clinical studies.

## Introduction

Oxygen-enhanced magnetic resonance imaging (OE-MRI) is a method that has been demonstrated for imaging lung function [1, 2]. Most OE-MRI studies to date have made use of T_1_-weighted acquisitions, which enable regional investigation of pulmonary ventilation and gas exchange across the alveolar epithelium into the bloodstream, since a change in T_1_ occurs due to the paramagnetic nature of oxygen dissolved in the parenchyma. During the last two decades, investigators have shown promising OE-MRI results for evaluating abnormal lung function in patients with chronic lung disease including interstitial pneumonia, pulmonary emphysema, cystic fibrosis, pulmonary thromboembolism, chronic obstructive pulmonary disease (COPD), lung cancer, and asthma [3–9].

To date, most OE-MRI studies in lungs have been performed at field strengths of 1.5 T or lower [10], while methodologies for lung OE-MRI have not been established at 3 T [11]. This is due to the inherent difficulties of lung MRI at higher field strengths; magnetic susceptibility differences at the numerous air–tissue interfaces within the lung increase and significantly shorten T_2_* in the parenchyma, thereby reducing the signal available for gradient echo-based methods. T_1_ relaxivity of oxygen decreases with increased field strength [12], which additionally hampers the sensitivity of the commonly-used ΔR_1_-based OE-MRI methods. Furthermore, if using gradient echo-based methods, the competing ΔR_2_* effect becomes substantial and dominates over the ΔR_1_ effect at 3 T, even at short TE [13]. Nevertheless, spoiled gradient echo pulse sequences are the most widely used methods for dynamic MRI data collection, enabling rapid acquisition of images with good spatial coverage and resolution. Although spin echo based- and ultrafast echo based-methods are not compromised by T_2_* effects, they may be limited to T_1_-sensitised ‘static’ (e.g. breath-hold or gated) OE-MRI due to relatively low temporal resolution. When considering the increasing availability of 3 T MRI, the above technical challenges motivate the developments of new methods in order to enable broad deployment of dynamic OE-MRI [11].

In addition, for OE-MRI to be used in clinical research and, eventually in clinical practice, it is essential to harmonise MRI protocols across centres and vendors. This requires a comparison of protocols, and selection and validation of comparable vendor-specific sequences as well as parameters. Derived biomarkers must demonstrate acceptable repeatability and reproducibility [14]. The purposes of our study were therefore 1) to use simulations to characterise the OE-MRI signal across a range of achievable sequence parameters at 3 T; 2) to evaluate the feasibility of a T_2_*-sensitized OE-MRI method at 3 T in healthy volunteers; and 3) to assess the scan-rescan repeatability and reproducibility of percent signal enhancement (PSE) and mean ΔR_2_* derived from dynamic OE-MRI in healthy volunteers across two sites and two vendors.

## Methods

For dynamic multi-slice OE-MRI acquisition, we implemented a dual echo RF-spoiled gradient echo sequence to enable estimation of T_2_*. We aimed to obtain images with a high temporal resolution to minimise motion artefact during free-breathing while maximising lung coverage and enabling reasonable spatial resolution. We determined that TR = 16 ms and matrix size = 96 × 96, would enable dynamic temporal resolution < 2 s and acquisition of six slices. Both TEs for the dual echo acquisition should be as short as possible to avoid losing signal due to low T_2_* and flip angle (FA) should be chosen to maximise signal difference between normoxia and hyperoxia.

### Simulations

We simulated the signal behaviour of the dual echo RF-spoiled gradient echo sequence at our chosen TR (16 ms) over a range of FA and TE values, described as follows [15]. First, the expected signal difference between air breathing and 100% oxygen breathing (ΔS) and PSE (100% x ΔS/S(air)) in the lung was simulated as a function of FA up to 30° and TE up to 3 ms using the following parameters: T_1_ (air) = 1281 ms [16], T_1_ (100% O_2_) = 1102 ms [16], T_2_* (air) = 0.68 ms and T_2_* (100% O_2_) = 0.62 ms. T_2_* values used in the simulations were calculated by averaging T_2_* values across all subjects’ values within this study (described below) as we were unable to identify previously-reported lung hyperoxic T_2_* values in the literature. Then, the expected ΔS values were simulated as a function of TE using FA of 5°, which was shown to maximise the absolute signal difference between the 21 % and 100 % oxygen images. Simulations were performed in MATLAB R2022b (MathWorks, Natick, MA).

### Participants & MRI Acquisition

Following approval of the local research ethics committee (REC Ref 18837/001) and written informed consent, we recruited 16 healthy volunteers with no previous record of lung diseases. Of these, eight subjects (3 males, age range = 26-54 years, median = 39.5 years) underwent lung MRI on a 3 T whole-body scanner from two vendors in different cities (Philips Ingenia in London, UK and Siemens MAGNETOM Vida in Manchester, UK) at a 4 week interval. Eight separate subjects (4 males, age range = 23-51 years, median = 27) were scanned twice to assess scan-rescan repeatability at a 4-6 week interval using the Philips scanner. Where possible, protocols for the two different scanner manufacturers utilised identical acquisition parameters, while some options required manufacturer-specific parameters. Site-independent parameters included: six coronal slices of 10 mm thickness with 4 mm gap with phase encoding Right/Left; in-plane resolution 4.69 x 4.69 mm^2^; field-of-view (FOV) covering entire lungs in all 16 volunteers (except for the inter-slice gaps in the anterior/posterior direction); TR = 16 ms; matrix size = 96 × 96; and dynamic temporal resolution = 1.54 s. Selection of FA (5°) was based on our simulations (Fig. 1). The shortest TE values available for the chosen acquisition were selected for each scanner (L, Philips in London; M, Siemens in Manchester): TE_1L_ = 0.71 ms, TE_2L_ = 1.2 ms; TE_1M_ = 0.81 ms, TE_2M_ = 1.51 ms. Scan parameters for each vendor are listed in Table 1.

**Figure 1.**
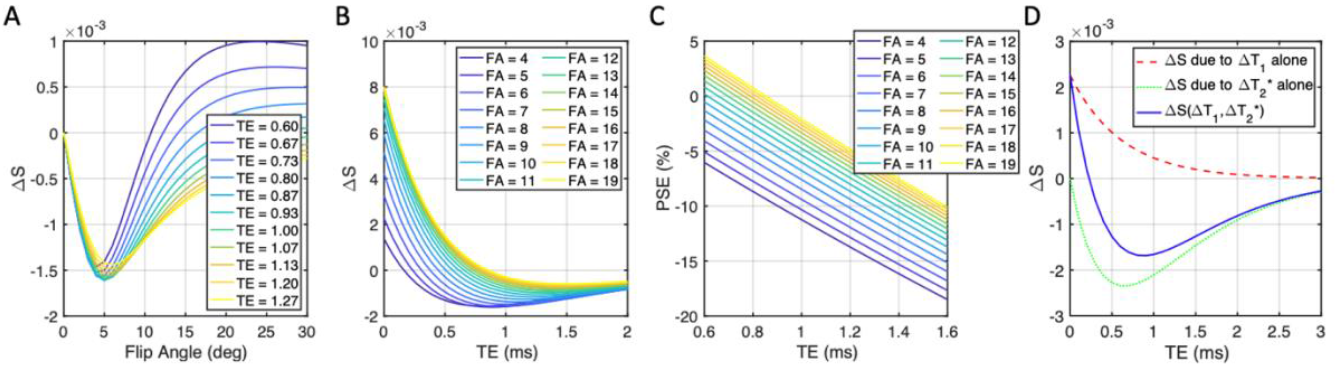
(A) The predicted OE signal change ΔS plotted as a function of flip angle at multiple TE, with TR = 16 ms. (B) The predicted signal change ΔS plotted as a function of TE with multiple flip angle, with TR = 16 ms. (C) Percent signal change (PSE) plotted as a function of TE at multiple flip angles, with TR = 16 ms. (D) The expected OE signal change for the T_1_-weighted RF spoiled gradient echo acquisition at 3 T due to ΔT_1_ alone (red dashed line), ΔT_2_* alone (green dotted line), and both (blue solid line), assuming literature-reported values for T_1_ and measured T_2_* in the lungs at 21% oxygen and 100% oxygen, flip angle = 5°, and TR = 16 ms.

**Table 1.** Scan parameters

|  | Scanner in London | Scanner in Manchester |
| --- | --- | --- |
| Manufacturer | Philips | Siemens |
| Model | Ingenia | MAGNETOM Vida |
| Field strength (T) | 3.0 | 2.9 |
| RF coil used | 32-channel torso coil in combination with the posterior coil | 18-channel body coil in combination with the 32-channel spine coil |
| Max. gradient strength (mT/m) | 45 | 45 |
| Max. slew rate (T/m/s) | 200 | 200 |
| TR (ms) | 16 | 16 |
| Echoes | full | half |
| Minimum achievable TE<sub>1</sub> (ms) | 0.71 | 0.81 |
| Minimum achievable TE<sub>2</sub> (ms) | 1.2 | 1.51 |
| FOV (mm x mm) | 450 x 450 | 450 x 450 |
| No of slices | 6 | 6 |
| Slice thickness (mm) | 10 | 10 |
| Gap (mm) | 4 | 4 |
| Acquired matrix | 96 x 96 | 96 x 96 |
| Orientation | Coronal | Coronal |
| Pixel size (mm x mm) | 4.7 x 4.7 | 4.7 x 4.7 |
| Flip Angle (°) | 5 | 5 |
| Bandwidth (Hz/Px) | 4488 | 2000 |
| Parallel Imaging | N | N |
| NSA | 1 | 1 |
| Time resolution (s) | 1.54 | 1.54 |
| Number of dynamics | 340 | 340 |

Subjects were fitted with a disposable/MRI-compatible non-rebreathing mask (Intersurgical, Berkshire, UK) to allow for medical air and 100% oxygen delivery while lying supine in the scanner. Piped gases were delivered to the subject at 15 l/min using a standard low flow oxygen blender (Inspiration Healthcare, Leicestershire, UK). The initial 60 dynamic acquisitions were obtained while breathing medical air. The gas supply was then switched to 100% O_2_ for the following 150 dynamic acquisitions, after which the supply was returned to medical air for further 130 acquisitions. Images were acquired during free-breathing. Total scanning time for the dynamic series was approximately 9 min.

### Data Analysis

After non-linear image registration of the free-breathing time series for motion correction using Advanced Normalization Tools (ANTs) [17, 18], the lung parenchyma, excluding central major vasculature, was manually segmented from registered images. For an initial exploration of the data, mean PSE maps were calculated by the subtraction of averaged hyperoxia images (61^st^ to 210^th^) from averaged normoxia images (10^th^ to 60^th^), normalized to the averaged normoxia images, following image registration and density correction as described below.

For our main data analysis, the dynamic series were fitted using exponential functions to characterize oxygen wash-in (encompassing the downslope and plateau regions of the curve) and wash-out (the upslope and return to baseline). The baseline for the exponential fit was defined as the averaged signal intensity across all normoxia time points before O_2_ inhalation as described in Equation 1 and the two functional forms were fitted to the curve indicated in Equations 2 and 3 for the downslope and upslope, respectively,

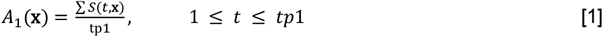

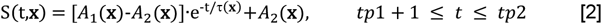

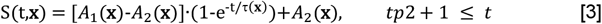

where A_1_(**x**) and A_2_(**x**) are the baseline and fitted negative maximum hyperoxia intensity (or plateau value) at position **x**, respectively, and τ, tp1 and tp2 are the fitted wash-in time constant and the provided gas switching time points (i.e. tp1, air to O_2_; tp2, O_2_ to air), respectively. Maximum PSE maps were produced by the subtraction of the baseline from the negative maximum hyperoxia value (A_1_-A_2_), normalized to the baseline A_1_.

As differences in lung tissue density can influence the measured signal enhancement between normoxia and hyperoxia, time-varying PSE maps were calculated twice, with and without a voxel-wise tissue density correction. Uncorrected PSE values were calculated by the subtraction of normoxia signal (S_21_) from hyperoxia signal (S_100_), normalized to S_21_ as

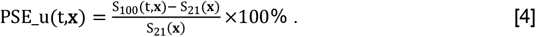

Tissue density variation was corrected using the adapted sponge model [19]. The whole-lung fractional volume change V was calculated at each time point by averaging the Jacobian determinant from the registration over all voxels in the defined lung mask across all slices. The respiratory index α_local_ was estimated voxel-wise (locally at the position **x** by linear regression estimation of the observed signal intensity S as a function of V as

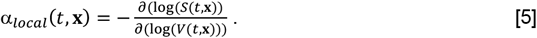

Then, the α_local_ values were applied as a voxel-wise density correction as

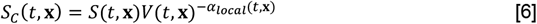

where S_C_(t,**x**) represent the corrected S(t,**x**). Corrected PSE values were quantified as

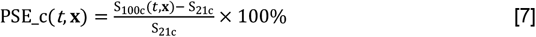

where S_100c_(*t***,x**) and S_21c_(*t*,**x**) represent the corrected S_100_(*t*,**x**) and S_21_(*t*,**x**), respectively. For comparisons between the pre- and post-density correction and for inter-scanner repeatability assessment, median PSE values within masks were calculated at each TE over the 2 most posterior slices for TE_1L_, TE_1M_, TE_2L_ and TE_2M_; anterior slices were excluded from the analysis due to poor signal to noise ratio (SNR) at TE_1M_ and TE_2M_. The median PSE value for each slice was then averaged across all slices and subjects. For intra-scanner scan-re-scan repeatability, median PSE values were averaged over 6 slices for TE_1L_ and TE_2L_, separately.

The R_2_* of each voxel was quantified analytically from the magnitude-reconstructed signal from the masked lung images acquired at TE_1_ and TE_2_ after tissue density correction as described in Eqs 5 and 6. ΔR_2_* maps were calculated by the subtraction of mean normoxia R_2_* maps across multiple time points (30^th^ to 60^th^ time series acquisitions) from mean hyperoxia R_2_* maps across multiple time points (120^th^ to 180^th^). Median ΔR_2_* values were averaged over two most posterior slices for multi-site comparison between Manchester and London, and six slices for scan-rescan comparison in London. Data analysis was performed in MATLAB R2022b (MathWorks, Natick, MA).

### Statistical Analysis

In the scan-rescan analysis of repeatability, we used Bland-Altman plots with 95% limits of agreement (LOA) and derived the repeatability coefficient (RC), reproducibility coefficient (RDC), and intraclass correlation coefficient (ICC) to evaluate the agreement of repeated measures [20]. The following levels of agreement were used: excellent for ICC > 0.74, good for ICC 0.6–0.74, fair for ICC 0.4–0.59 and poor for ICC < 0.4 [21]. All statistical analyses were performed using SPSS v28.0 (SPSS Inc, Chicago, IL).

## Results

### The effect of oxygen on signal intensity – simulations

Simulations show that ΔT_2_*-induced negative enhancement (ΔS) for our chosen TR = 16 ms is maximum at FA ~ 5°, largely independent of the choice of TE (Fig. 1A). The amplitude of the TE dependence of ΔS reduces with smaller FA (Fig. 1B), with negative-going signal change occurring at shorter TE. The magnitude of negative PSE increases with lower FA and longer TE, while PSE values are closer to 0 at shorter TE and high FA (Fig. 1C). We also observed that ΔT_2_* dominates the signal change and produces negative contrast at TE > 0.2 ms for FA = 5° (Fig. 1D). The expected signal change at TE_1L_ (0.71 ms) is about 55 % more sensitive to changes in ΔT_1_ and about 21 % more sensitive to changes in ΔT_2_* than at TE_2L_ (1.2 ms) (Fig. 1D).

### The effect of oxygen on signal intensity – experimental

Typical location of the acquired images is shown in Fig. 2A. PSE maps at both TEs demonstrate uniform PSE across the parenchyma (Figs. 2B, 2D). As expected, due to the dominant effect of T_2_* changes, time course plots of mean PSE from masked lungs exhibit negative contrast due to 100% O_2_ inhalation (Figs. 2C, 2E), in agreement with the simulated results (Fig. 1D). The mean signal intensity is higher in posterior slices due to greater proton density, associated with the subjects’ supine position (Figs. 2C, 2E, supporting information Fig. S1).

**Figure 2.**
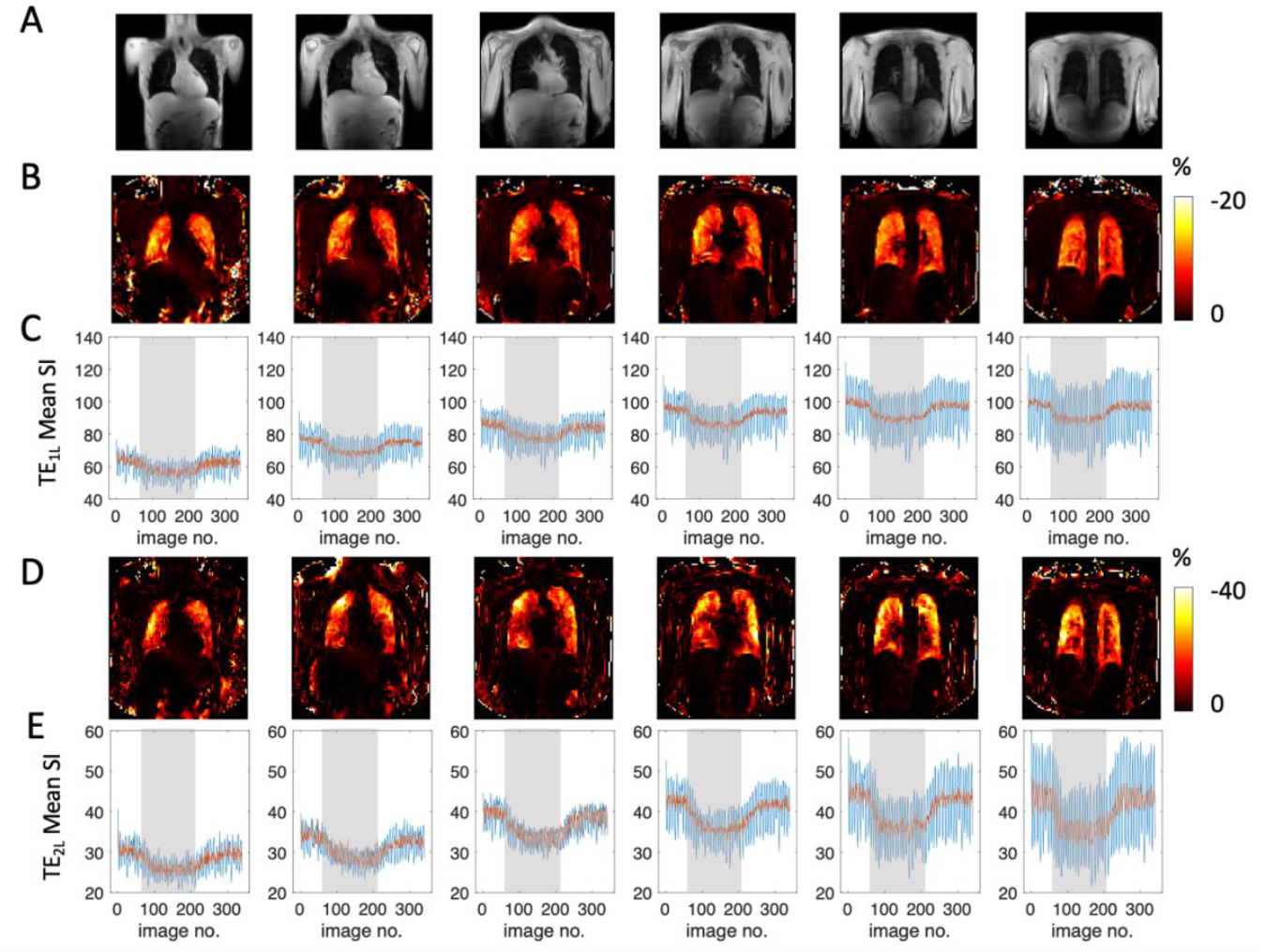
(A) The typical location of the six slices from anterior to posterior. Example subject data show unmasked percentage signal change maps obtained with (B) TE_1L_ (0.71 ms) and (D) TE_2L_ (1.2 ms), and (C and E) the corresponding time course curves of the mean signal intensity from masked, registered lung for each slice. Blue lines show uncorrected signal; red line shows signal after density correction.

### The effect of density correction

The time course plots (Figs. 2C, 2E) post-density correction (red solid line) show smaller magnitude signal fluctuation than pre-density correction (blue solid line) due to the reduced impact of respiratory motion-induced signal changes. Example signal time courses with their downslope and upslope fits (Eqs. 2, 3) also show improvement in time course wash-in fitting with tissue density correction (supporting information Fig. S2).

Table 2 summarises the mean PSE and coefficient of variation (CV) pre- and post-tissue density correction. Across eight healthy volunteers, the magnitudes of all negative PSE values were reduced by applying tissue density correction. In addition, CV of mean PSE was also decreased for all except TE_1M_ PSE. All repeatability metrics were improved by the density correction step (supporting information Fig. S3) as also described in section on ‘Repeatability and Reproducibility’.

**Table 2.** Inter-subject reproducibility and intra-scanner repeatability assessment of uncorrected and corrected mean (± standard deviation (std)) fitted PSE measurements from eight travelling healthy volunteers in four different TEs (TE_1L_ and TE_2L_ in London; TE_1M_ and TE_2M_ in Manchester). The mean PSE and CV were calculated across all subjects at each TE based on the measurement from the two most posterior slices. Difference (D), the Bland-Altman (BA) 95% limit-of-agreement (LOA), repeatability coefficient (RC) and intra-scanner variation (ICC_intra_) were calculated from all six slices between two scan-rescan sessions in London for each subject.

| | | TE (ms) | Mean PSE (%) ± std (%) | CV (%) | Difference (%) | 95% LOA (%) | RC (%) | $ICC_{intra}$ |
| --- | --- | --- | --- | --- | --- | --- | --- | --- |
| Uncorrected PSE mapping (Eq 4) | $TE_{1L}$ PSE | 0.71 | -9.61 ± 4.24 | 44.13 | -1.33 | (-7.22, 4.55) | 8.32 | 0.05 |
| | $TE_{1M}$ PSE | 0.81 | - 8.61 ± 1.97 | 22.84 | | | | |
| | $TE_{2L}$ PSE | 1.20 | - 14.87 ± 4.51 | 30.35 | -0.67 | (-7.53, 6.20) | 9.71 | 0.23 |
| | $TE_{2M}$ PSE | 1.51 | - 14.01 ± 2.77 | 19.80 | | | | |
| Corrected PSE mapping (Eq 7) | $TE_{1L}$ PSE | 0.71 | -6.55± 1.92 | 29.32 | -0.51 | (-2.36, 1.33) | 2.61 | 0.77 |
| | $TE_{1M}$ PSE | 0.81 | -8.06 ± 1.84 | 22.85 | | | | |
| | $TE_{2L}$ PSE | 1.20 | -12.20 ± 2.75 | 22.56 | -0.57 | (-2.99, 1.84) | 3.42 | 0.92 |
| | $TE_{2M}$ PSE | 1.51 | -13.37 ± 2.46 | 18.39 | | | | |

### Echo time dependence

While the mean signal intensity is higher at TE_1L_ than for TE_2L_ (Figs. 2C, 2E), PSE is greater at TE_2L_ than at TE_1L_ (Figs. 2B, 2D), again in agreement with our simulations (Figs. 1B, 1C). The TE dependence of PSE expected from simulations (Fig. 1C) is also observed in the density-corrected PSE values from four separate TEs at two sites (Table 2).

### ΔR_2_* quantification

Figure 3 shows examples of plateau ΔR_2_* maps across six slices from anterior to posterior, the corresponding time course plots of the median R_2_* from the maps of masked lungs for each slice. Median ΔR_2_* maps illustrate clear O_2_ delivery in the entire lung (Fig. 3A, 3C), with a spatial distribution that is less homogeneous than seen in the PSE maps (Fig. 2). Median R_2_* time course plots show R_2_* is largely unaffected by density correction (because the calculation of R_2_* normalises for density) except for the posterior slice, which is affected by partial volume (Fig. 3B). A trend of ΔR_2_* increase from anterior to posterior slices is observed (Fig. 4C). The mean T_2_* values of sixteen healthy volunteers from two posterior slices were 0.68 ± 0.05 ms (T_2_* for 21% O_2_), 0.62 ± 0.05 ms (T_2_* for 100% O_2_) and mean ΔR_2_* was 0.14 ± 0.03 ms^-1^ (Table 3).

**Figure 3.**
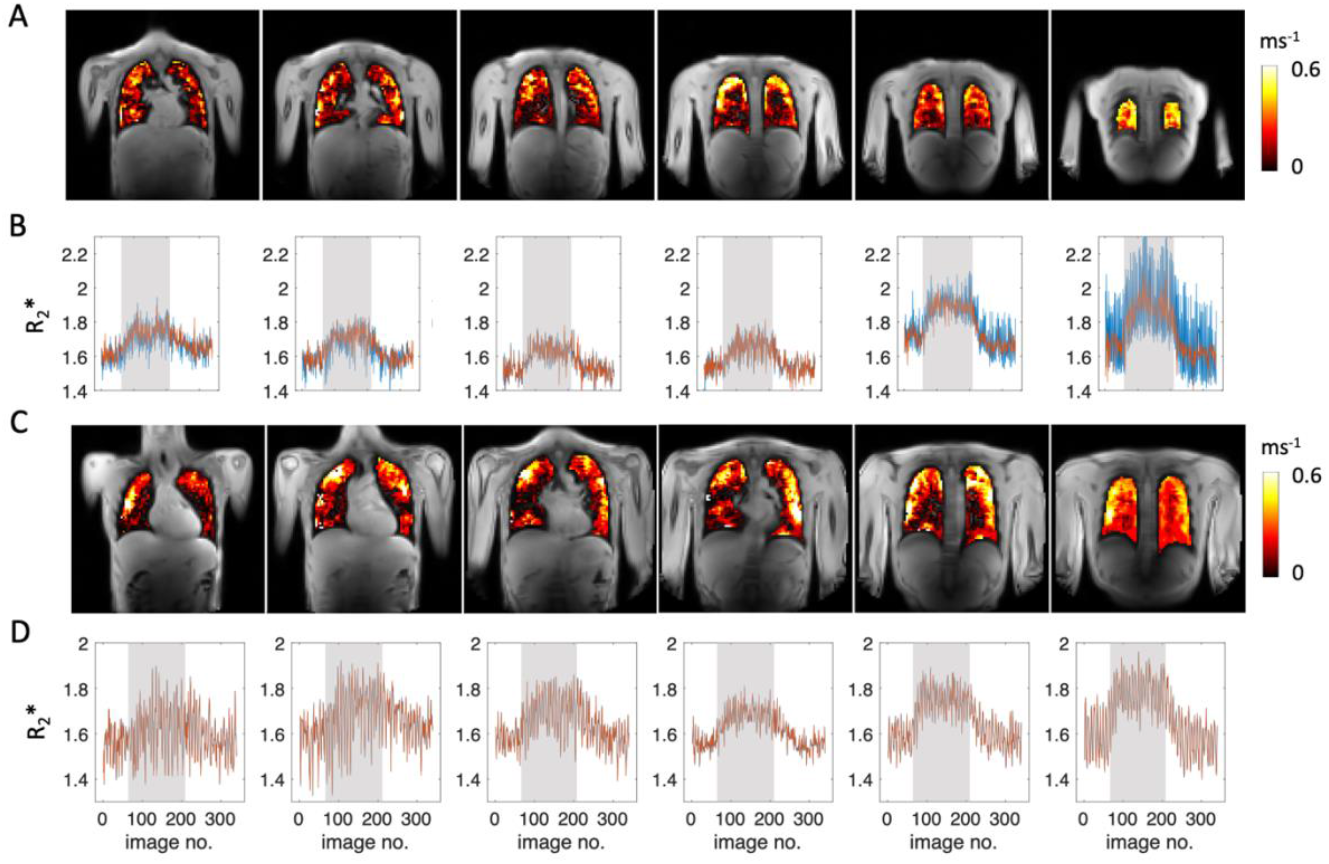
Examples of two subjects: (A, C) the plateau ΔR_2_* maps of masked lung, six slices from anterior to posterior and (B, D) the corresponding time course curves of median R_2_* from masked lung along for each slice. Increase of R_2_* due to 100% O_2_ inhalation is visible in all slices but clearer in posterior slices.

**Figure 4.**
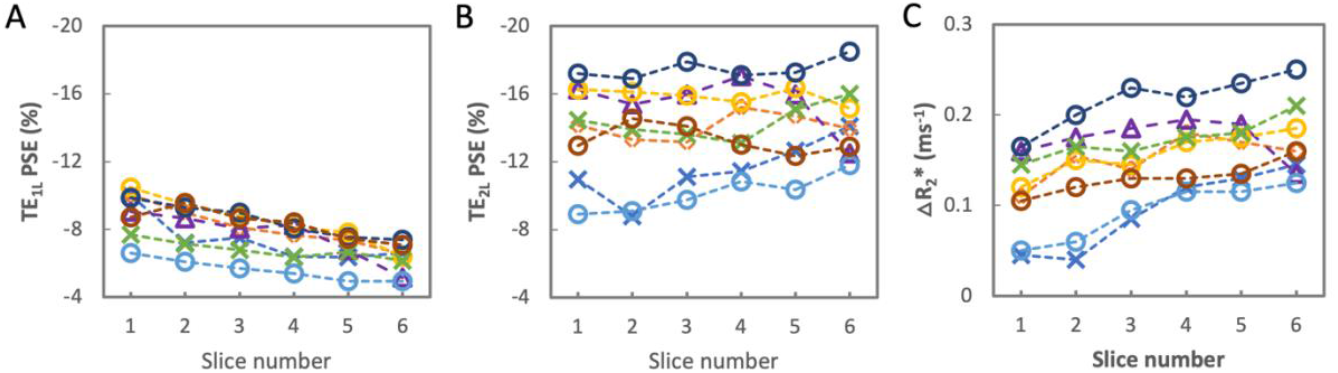
The PSE from masked, registered, tissue density corrected lung for each slice of eight individual subjects scanned in London with (A) TE_1L_ (0.71 ms) and (B) TE_2L_ (1.2 ms). (C) Equivalent plot for ΔR_2_*.

**Table 3.**
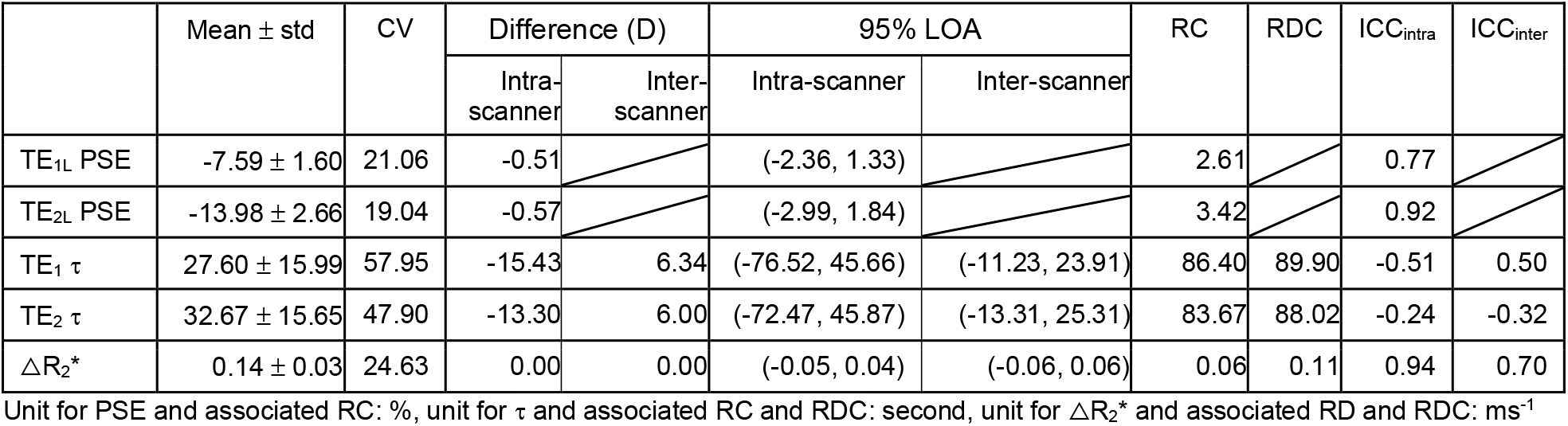
Repeatability and reproducibility for percentage signal enhancement (PSE) and ΔR_2_*. First and second columns: mean ± standard deviation (std) and coefficient of variation (CV) of PSE from the two repeated sessions in London, and wash-in time (τ) values and ΔR_2_* from travelling volunteers at both sites and the two repeated sessions in London. Middle columns: mean difference between two sessions (D), the Bland-Altman (BA) 95% limit-of-agreement (LOA) for inter- and intra-scanner comparisons, repeatability coefficient (RC) for intra-scanner comparisons and reproducibility coefficient (RDC) for inter-scanner comparisons. Last two columns: intra-class correlation coefficient (ICC) for intra-scanner variation (ICC_intra_) and for inter-scanner variation (ICC_inter_) based on absolute agreement, 2-way mixed-effects model.

### Signal variation with slice position in the lung

The fitted PSE of TE_1L_ gradually decreases from anterior to posterior slices across all subjects (Fig. 4A) whereas the PSE of TE_2L_ do not noticeably change (Fig. 4B). ΔR_2_* shows a gradual increase from anterior to posterior.

### Repeatability and Reproducibility

Example fitted PSE maps from an intra-scanner, intra-subject scan-rescan of a healthy volunteer show relatively homogeneous enhancement at both TEs (Figs. 5A, 5B). The mean fitted PSE values from eight healthy volunteers varied little between the repeat scans (−7.39 % ± 1.61 % and −7.79 % ± 1.58 % at TE_1L_; −13.71 % ± 2.96 % and −14.27 % ± 2.31 % at TE_2L_). The mean fitted PSE values across both sessions were −7.59 % and −13.98 % at TE_1L_ and TE_2L_, respectively (Table 3). The mean wash-in time (τ) values across all subjects were 27.60 s and 32.67 s at TE_1_ and TE_2_, respectively (Table 3). Example wash-in time maps from an intra-scanner, intra-subject scan-rescan of a healthy volunteer and inter-scanner scans of a travelling volunteer show low repeatability and reproducibility at both TEs (supporting information Fig. S4).

**Figure 5.**
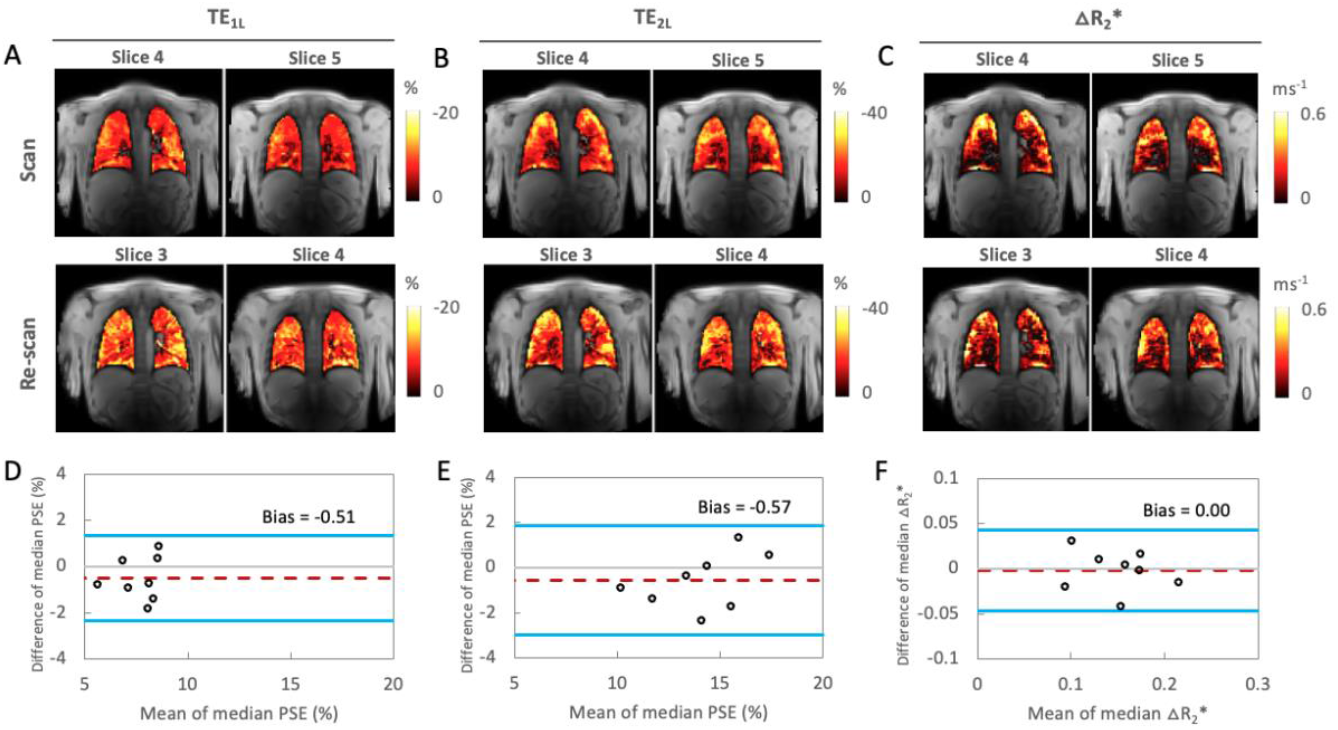
Bland-Altman analysis comparing PSE and ΔR_2_* between two separate sessions (repeatability) in London. (A) mean PSE with TE = 0.71 ms, (B) mean PSE with TE = 1.2 ms, (C) mean ΔR_2_*, and (D and E) Bland-Altman plots for the repeated measurements of PSE from the 1^st^ and 2^nd^ TE and (F) ΔR_2_* (intra-scanner, intra-subject).

The plateau ΔR_2_* maps from the same data set show relatively heterogeneous ΔR_2_* distribution, with some areas in areas of major vasculature that do not appear to respond to 100% O_2_ inhalation (Fig. 5C). This is expected as the R_2_* change is mainly due to gaseous oxygen in the alveoli but not dissolved oxygen as previously reported [13].

The Bland-Altman plot analysis of the repeated measurements of PSE and ΔR_2_* (intra-scanner, intra-subject) is shown in Figs. 5D–5F. Little suggestion of bias between measurements is evident, with the 95% LOA for the two repeated PSE measurements in London being [-2.36 %, 1.33 %] at TE_1L_ (0.71 ms) and [−2.99 %, 1.84 %] at TE_2L_ (1.2 ms) (Figs. 5D, 5E). For ΔR_2_* between two separate sessions, the 95% LOA measurements were [−0.05 %, 0.04 %] (Fig. 5F). The ICC and RD measurements of fitted PSE at TE_1L_ and TE_2L_, and ΔR_2_* for intra-scanner, intra-subject (ICC_intra_) were 0.77 (RD = 2.61 %), 0.92 (RD = 3.42 %) and 0.94 (RD = 0.06 ms^-1^), respectively, suggesting excellent scan-rescan repeatability (Table 3). Wash-in time τ showed low scan-rescan repeatability at both TEs, with ICC_intra_ of −0.51 (TE_1L_) and −0.24 (TE_2L_) and RC of 86.40 s (TE_1L_) and 83.67 s (TE_2L_).

The fitted PSE maps from inter-scanner travelling volunteers repeat scans show similar spatial distribution of enhancement at TE_1_ while the enhancement at TE_2_ from the Manchester site is more noisy, potentially due to signal approaching the noise floor at the longer TE (Fig. 6A). Plots of the combined fitted PSE values at 4 separate TEs from the two MRI systems display the expected TE dependence of the signal (Fig. 6B, Table 2), similar to simulation (Fig. 1C, supporting information Fig. S5). The ΔR_2_* maps again show a lack of ΔR_2_* enhancement near major vasculature (Fig. 6C). The ΔR_2_* maps from the Manchester site are again more noisy, as observed with the TE_2_ data. However, ΔR_2_* comparison between two different scanners again shows little evidence of bias, with the 95% LOA measurements being [-0.06 %, 0.06 %] (Fig. 6D). Good ICC and RDC values were observed for inter-scanner ΔR_2_* comparisons (ICC_inter_ = 0.70; RDC = 0.11 ms^-1^, Table 2). Wash-in time from the inter-scanner travelling volunteers repeat scans show low ICC and high RDC values at both TEs, indicating poor reproducibility (Table 2).

**Figure 6.**
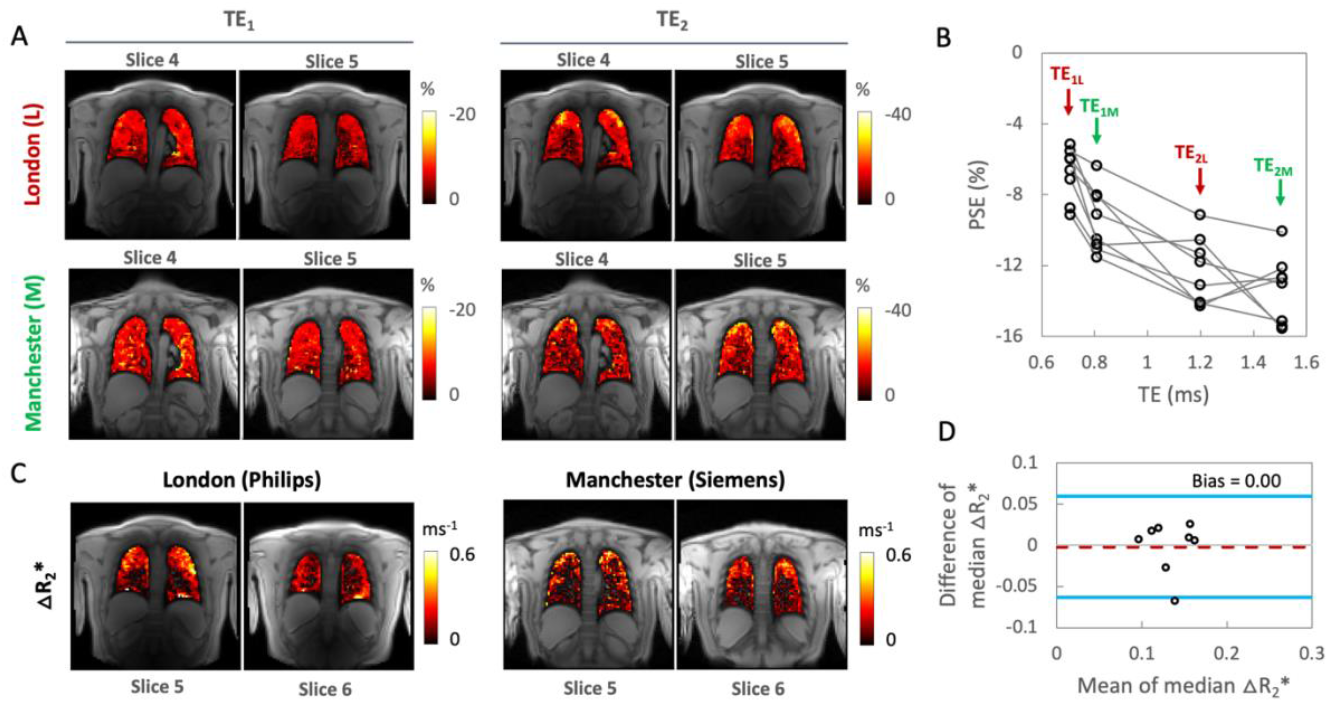
An example of inter-scanner intra-subject reproducibility of PSE at (A) TE_1_ and TE_2_ scanned using two MRI systems at two sites (London and Manchester). (B) PSE obtained at 4 separate echo times (TE_1L_ = 0.71 ms, TE_2L_ = 1.2 ms in London and TE_1M_ = 0.81 ms, TE_2M_ = 1.51 ms in Manchester). The combined PSE values from two MRI systems show a similar trend as a function of TE as the PSE simulation (Fig. 1C). (C) Inter-scanner intra-subject reproducibility of ΔR_2_* from the same subject. (D) Bland-Altman analysis comparing ΔR_2_* between two scanners for the same subjects.

## Discussion

In recent years, installations of a clinical 3 T MR systems have significantly increased worldwide, often motivated by the higher SNR that can frequently be achieved, relative to lower field systems. However, the viability of dynamic OE-MRI at 3 T has not to date been investigated. In this work, we demonstrate the feasibility of detecting dynamic OE signal change and of quantifying ΔR_2_* due to oxygen breathing at 3 T. To progress the translation of these biomarkers towards clinical use, we also evaluate the intra-scanner repeatability and the inter-scanner/cross-site reproducibility of the proposed method. Although two studies have reported the feasibility of 3 T T_1_-weighted OE-MRI and compared it with either 1.5 T MRI or CT [11, 22], to the best of our knowledge, this is the first report investigating detailed dynamic signal enhancement behaviour, repeatability and reproducibility, simultaneously with quantification of dynamic ΔR_2_* due to oxygen inhalation, using a dual echo RF-spoiled gradient echo acquisition at 3 T.

Our motivation for focussing on T_2_*-related contrast at 3 T is twofold. Firstly, the longitudinal relaxivity of O_2_ is approximately 20% lower at 3 T than 1.5 T [12], leading to a proportionately smaller achievable ΔR_1_ at 3 T. Secondly, T_2_* in lung decreases with field strength, meaning that SNR in T_1_-weighted OE-MRI is much reduced. T_2_*-based OE-MRI has been proposed to counter some of these detrimental effects, although previously-developed methods employed non-standard acquisition methods [23]. Of note, T_2_*-related signal is potentially more specific to ventilation as it is expected to be an effect of changing concentrations of oxygen gas in the alveoli rather than dissolved oxygen [13]. In this study, we optimised a multi-slice dual-echo RF-spoiled gradient echo acquisition. This method enables measurement of dynamic OE signal change at high temporal resolution with controllable T_2_*-weighting, and monitoring of dynamic ΔR_2_*, simultaneously, while requiring no or minimal pulse programming. This easy implementation on standard clinical platforms is intended to assist in clinical translation of this technique.

### T_2_*-weighting allows good oxygen delivery contrast at 3 T

Our simulations (Fig. 1D) show that spoiled gradient echo PSE is unsurprisingly dependent on both ΔT_1_ and ΔT_2_*, which can lead to reduced oxygen-related signal change if TE and flip angle are not optimised. Maximum (negative) PSE for our chosen TR of 16 ms is found with a flip angle of around 5° across a wide range of TE (Fig. 1A). Our simulations also indicate that negative PSE at TE longer than approximately 0.23 ms when using this flip angle is due to the significant oxygen-level dependent ΔR_2_* effect dominating the signal change in the lungs (Figs. 1B, 1D). Our experimental data are consistent with our simulations, with negative PSE observed throughout the lung parenchyma, at levels that are in agreement with simulations. Importantly, this allows the generation of visually high quality mean PSE maps at the TEs used in this study (Figs. 2, 5, 6). The mean fitted PSE values of travelling healthy volunteers from two sites/vendors ranged from −6.55 ± 1.92 % to −13.37 ± 2.46 % for TE = 0.71 ms to 1.51 ms (Table 2). These results are consistent with the expected trend of PSE with variable TEs from our simulations (Fig. 1C). The mean T_2_* value across all healthy volunteers for 21% O_2_ inhalation was 0.68 ± 0.05 ms, which is consistent with the reported values of 0.74 ± 0.1 ms at 3 T in literature [24]. We observed that upon 100% O_2_ inhalation, the mean T_2_* value decreased by about 9% relative to normoxia, which resulted in ΔR_2_* of 0.14 ± 0.03 ms^-1^. To the best of our knowledge, this is the first study that reports mean values of hyperoxic T_2_* and ΔR_2_* of healthy human lungs at 3 T.

While the influence of T_2_* on PSE is large at 3 T, the effect will also be present at lower field strengths when using gradient echo methods and should be accounted for when interpreting nominally T_1_-weighted OE-MRI [15]. Future work will explore the utility of our proposed double echo approach at lower field strengths.

### T_2_*-weighted OE-MRI demonstrates good repeatability and reproducibility

The Bland-Altman and ICC analysis of the repeated measurements of PSE and ΔR_2_* suggest high intra-scanner and intra-subject repeatability (Table 2, 3, Fig. 5). They also demonstrate that comparable dynamic OE-MRI protocols for the lung can be implemented at 3 T across different sites and scanners with good repeatability and reproducibility for ΔR_2_*. While maximum gradient strength and maximum gradient slew rate between the two systems from the different manufacturers are identical, matching TE and bandwidth between scanners proved challenging, which results in variability of OE signal enhancement. For this reason, we were unable to directly perform reproducibility assessments for PSE, although we did observe that the variation in PSE with TE between scanners matched our simulations well (Figs. 1C, 6B). We were able to assess ΔR_2_* reproducibility, as T_2_* signal decay is, to the best of our knowledge, monoexponential with TE in the lung. However, we observed significant variation in wash-in time, resulting in low intra-scanner repeatability and inter-scanner reproducibility of wash-in time. This may be attributed to inaccurate gas switching time points for the fitting as the gas blender is manually operated, or to differences in participants’ breathing patterns between scans, which have a direct impact on ventilation.

### Density correction improves repeatability

Our results demonstrate that the density correction based on the adapted sponge model significantly improves quantification of OE-MRI metrics by decreasing fluctuation due to respiratory motion-induced signal changes as (Table 2, supporting information Fig. S3). This is particularly useful in posterior slices (in supine position) where fluctuation of signal changes is greater (supporting information Figs. S1).

Our results also show that the proposed method at 3 T yields excellent intra-scanner repeatability after correction of pixel-wise signal intensity using the deformation fields from image registration (Table 3). Previous studies using non-Cartesian ultra-short TE (UTE) approach with free-breathing at 1.5 T [25] or breath-held acquisitions at 0.55 T [10] also demonstrated improvement of repeatability in both mean PSE and the low-enhancement percent by demonstrating reduced Bland-Altman limits of agreement and improved ICC when correction of pixel-wise signal intensity was applied.

### Signal variation with position in the lung

Mean signal intensity for both baseline and change due to 100 % oxygen is observed to be higher in posterior slices across all subjects due to greater proton density in subject’s supine position (Figs. 2C, 2E, supporting information Fig. S1). Interestingly, we also observed that the absolute PSE of TE_1_ gradually decreases, from anterior to posterior slices, across all subjects (Fig. 4A) whereas the PSE of TE_2_ does not noticeably change (Fig. 4B). While PSE combines both ΔT_1_ and ΔT_2_* effects, the PSE at TE_1_ contains a stronger ΔT_1_ effect than that at TE_2_, as shown in the simulation (Fig. 1D). On the contrary, ΔT_2_* effect is more substantial and dominating over ΔT_1_ effect in the PSE at TE_2_. Thus, this may be attributed to increasing effect of ΔT_1_ from anterior to posterior, possibly due to increased vessel density and/or blood pooling due to gravity, which require further investigation. A trend of ΔR_2_* increase observed from anterior to posterior slices may reflect the expected predominant sensitivity of ΔR_2_* to ventilation, as more ventilation is expected in the posterior slices when the lungs are in supine position. This is consistent with a previous report that T_2_*-related signal is potentially more specific to ventilation due to an effect of changing concentrations of oxygen gas in the alveoli [13].

A limitation to the present study is that we employed a 2D multi-slice readout, designed to ensure good temporal resolution. This allows good lung coverage, but slice gaps and the limited number of slices may mean that some localized pathology could be missed. This could be mitigated by increasing the number of slices, either by increasing TR (thereby lowering temporal resolution and leading to more motion-related image blurring and artefacts) or by employing acceleration methods. Since there may be inconsistencies between the slice positions, a multi-slice acquisition also leads to challenges for inter-scanner, inter-session image registration that is essential for voxel-wise comparison. 3D non-Cartesian UTE OE-MRI methods have been demonstrated at 1.5 T and lower field strengths to allow isotropic spatial resolution whole lung OE-MRI measurements [10, 15, 25]. However, those studies are limited to static acquisitions which employed either breath-hold or two separate free-breathing sessions of normoxia and hyperoxia. Dynamic OE-MRI using such methods may be possible by employing temporal view sharing methods, but we are unaware of any studies to date that have made use of this strategy.

Despite the above limitations, the present study demonstrates that our proposed dynamic OE-MRI method with transition between two gas phases enables us to robustly monitor oxygen wash-in and enhancement, which is likely to provide relevant functional information. Future studies in respiratory diseases will allow us to better understand this potential.

## Conclusion

Despite technical challenges of OE-MRI in the lung at higher magnetic field, we have shown the feasibility of lung dynamic OE-MRI at 3 T and signal enhancement behaviour in the lung using an optimized dual echo RF-spoiled gradient echo acquisition. The proposed method enables the simultaneous dynamic measurement of signal enhancement (PSE) and change in R_2_* at 3 T. Our results demonstrate excellent intra-scanner repeatability for PSE across time points and reproducible ΔR_2_* across two sites and vendors, suggesting potential utility in multi-centre clinical studies. As measuring PSE at cross-sites reveals a significant dependence of observed PSE on TE, any comparison of PSE values should also consider the TEs used.

## Acknowledgement

This work is supported by the Cancer Research UK National Cancer Imaging Translational Accelerator (NCITA) awards C1519/A28682 (UCL) and C19221/A28683 (University of Manchester), the EPSRC-funded UCL Centre for Doctoral Training in Medical Imaging (EP/L016478/1; EP/S021930/1), an EPSRC Industrial CASE award (Voucher No. V20000074), GlaxoSmithKline Research and Development Ltd (BIDS3000035683), and Innovate UK award 104629. We thank Lucy Caselton, Sumandeep Kaur and David M. Higgins for the technical assistance, and Stanley Kruger for the constructive discussion for the simulation.

