## Supporting Information for "Dynamic oxygen-enhanced MRI of the lung at 3 T: feasibility, repeatability and reproducibility"

Figure S1. Example time course curves of the median signal intensity (SI) and $R_2$* from masked, registered lung for each slice of a single travelling subject obtained in London (A) and Manchester (B) with $TE_{1L}$ (0.71 ms), $TE_{2L}$ (1.2 ms), $TE_{1M}$ (0.81 ms), and $TE_{2M}$ (1.51 ms), by pre- (blue line) and post-tissue density correction (red line).

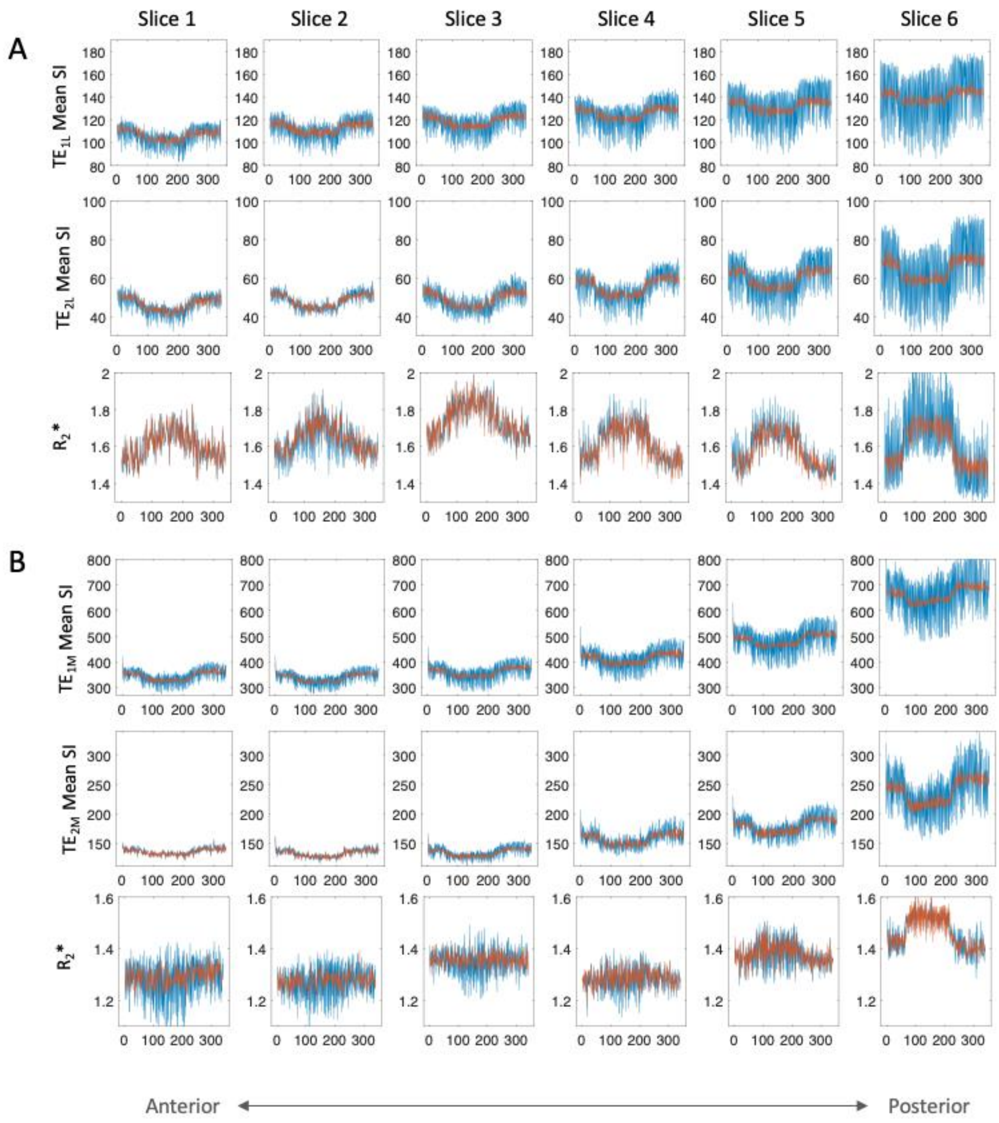

Figure S2. (A) Pre- and (B) post-density corrected example time course (blue dashed lines) and fits (red solid lines) for downslopes and upslopes from individual voxels.

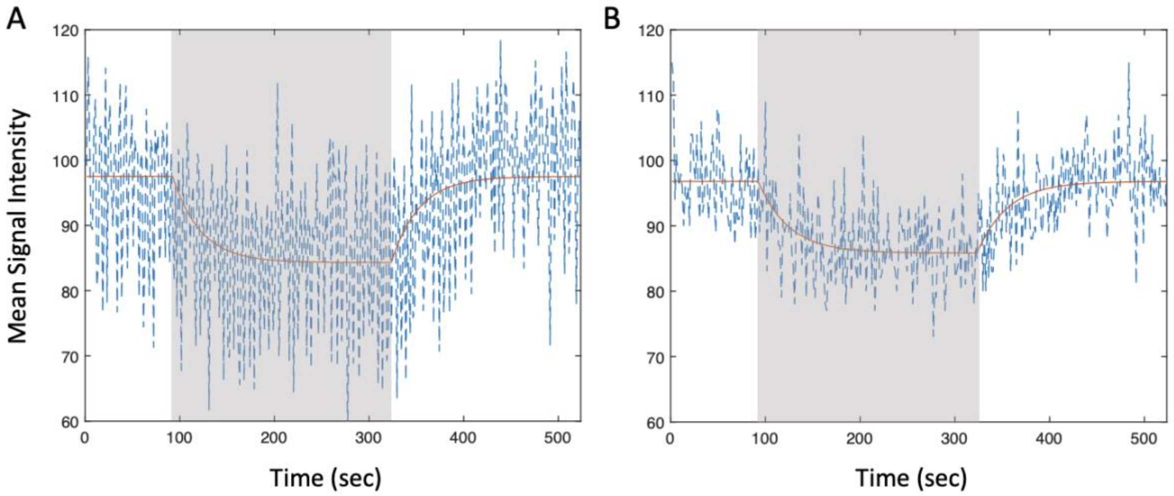

Figure S3. The Bland-Altman plots for the repeated measurements of percent signal change (PSE) from the 1st and 2nd TE before (A and C for $TE_{1L}$ and $TE_{2L}$, respectively) and after tissue density correction (B and D for $TE_{1L}$ and $TE_{2L}$, respectively). The 95% LOA decreased from [-7.22 %, 4.55 %] to [-2.36 %, 1.33 %] for $TE_{1L}$ and [-7.53 %, 6.20 %] to [-2.99 %, 1.84 %] for $TE_{2L}$. Similarly, additional statistical metrics display significantly reduced RC (69 % and 65 % for $TE_{1L}$ and $TE_{2L}$, respectively) and increased $ICC_{intra}$ (94 % and 75 % for $TE_{1L}$ and $TE_{2L}$, respectively) with tissue density correction compared to pre-density correction (see Table 2).

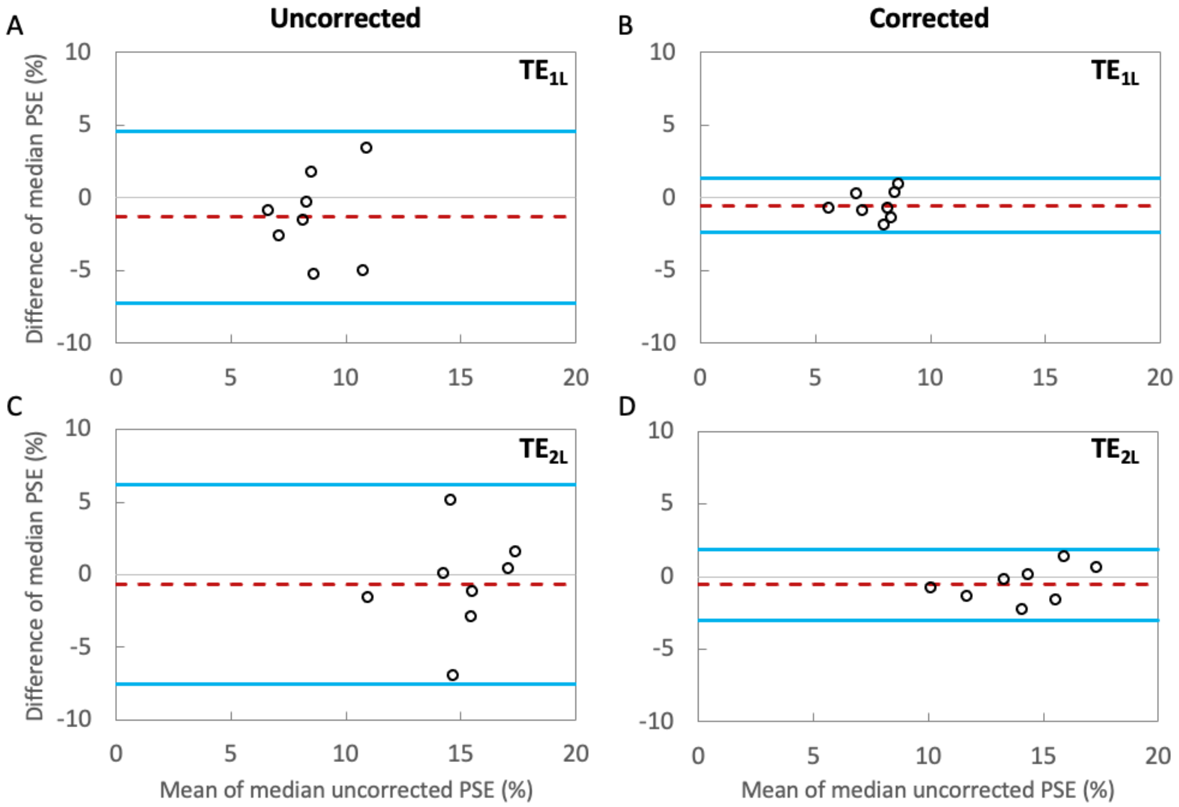

Figure S4. Examples of wash-in time maps of masked lung, six slices from anterior to posterior (A) between two separate sessions (repeatability) in London and (B) inter-scanner intra-subject reproducibility from a travelling volunteer

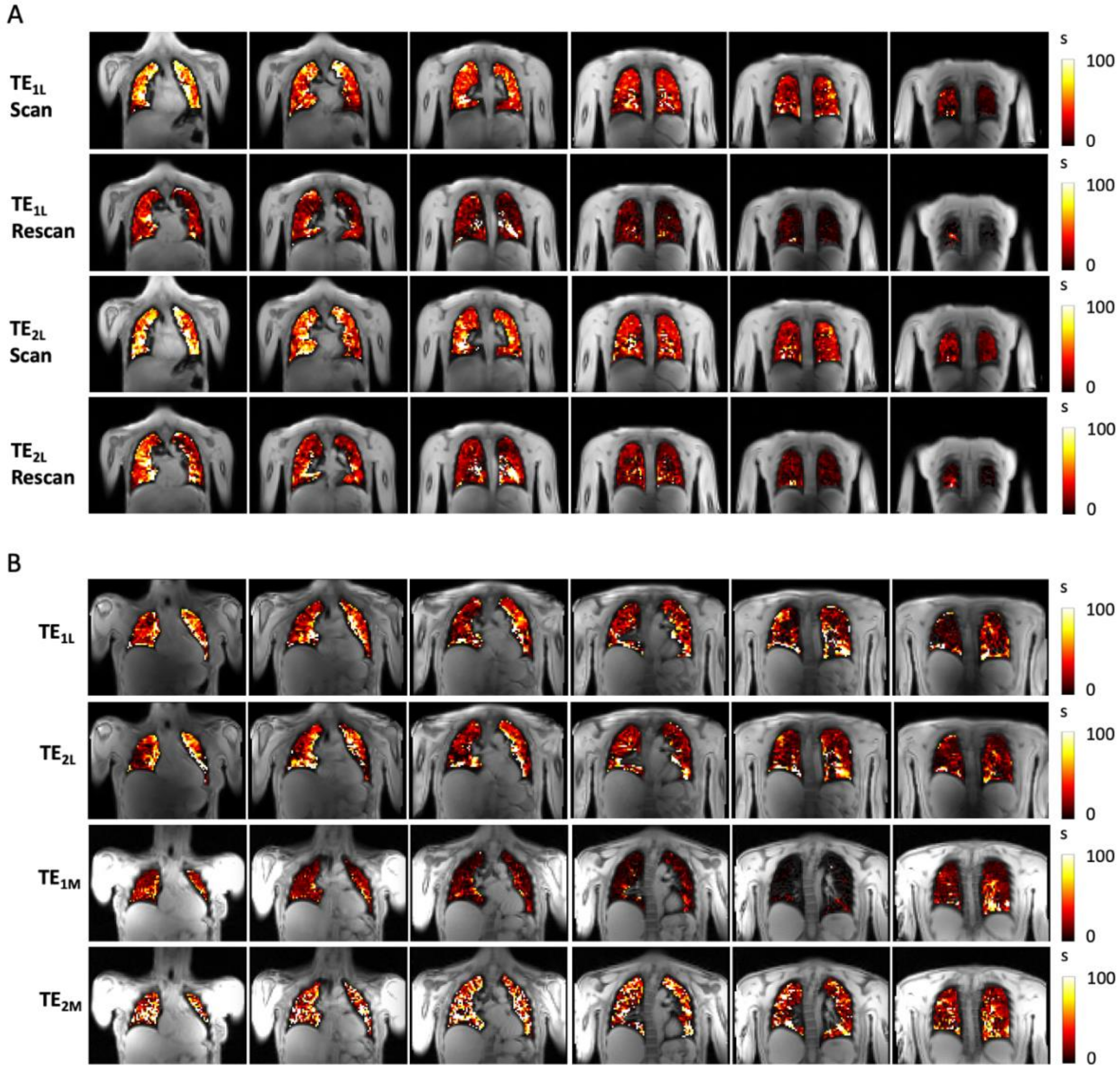

Figure S5. Simulated vs experimental percent signal change (PSE) values plotted as a function of TE, assuming literature-reported values for $T_1$ (1281 ms for air and 1102 ms for 100% $O_2$) and measured $T_2$* in the lungs at air and 100% oxygen, flip angle = 5°, and TR = 16 ms. The $T_2$* values were averaged over 8 travelling volunteers scanned in London (0.67 ms for air and 0.60 ms for 100% $O_2$). The experimental PSE values were obtained at 4 separate echo times ($TE_{1L}$ = 0.71 ms, $TE_{2L}$ = 1.2 ms in London and $TE_{1M}$ = 0.81 ms, $TE_{2M}$ = 1.51 ms in Manchester) and averaged over 8 travelling volunteers scanned in London and Manchester (as displayed in Fig. 6B of the main paper). The combined PSE values from two MRI systems show a similar trend as a function of TE as the PSE simulation with variability in longer TE.

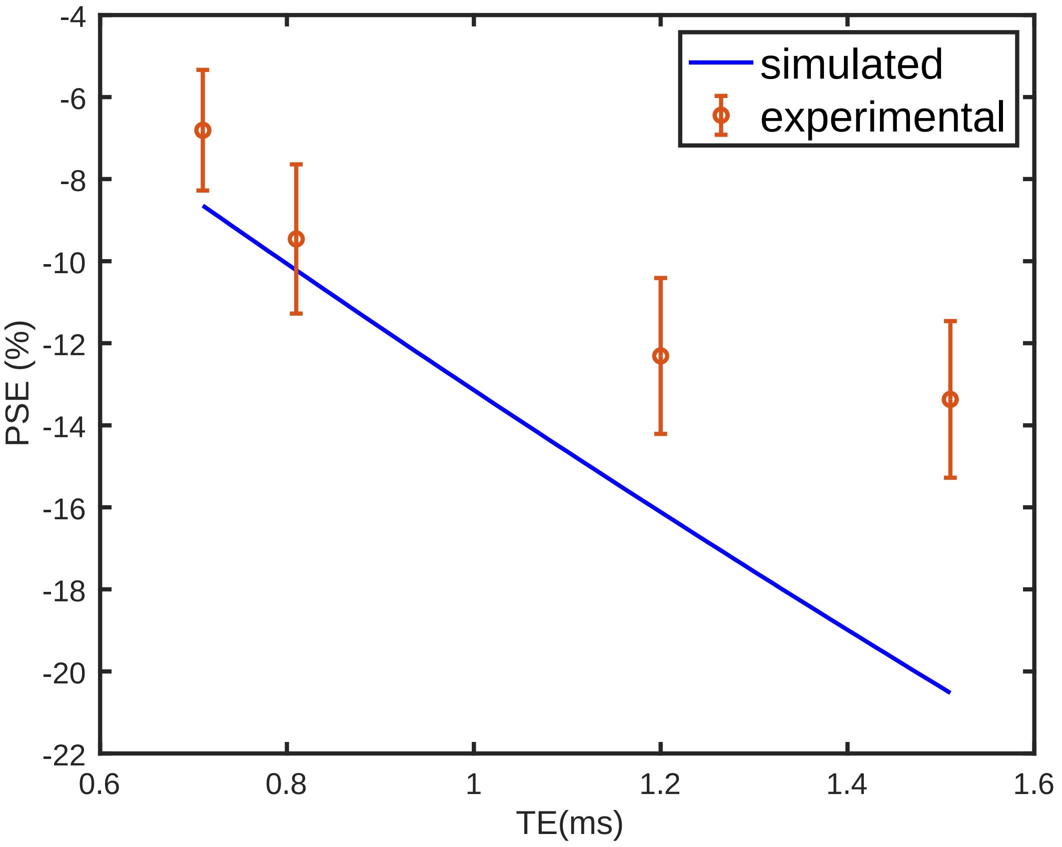
